# Heat stress reveals divergent pollen viability responses across species and populations of monkeyflower

**DOI:** 10.64898/2026.09.25.754521

**Authors:** Derek A. Denney, Caitlin N. Byron, David B. Lowry

## Abstract

**Premise:** Rising temperatures can reduce plant reproductive success by disrupting pollen development, yet little is known about how reproductive heat sensitivity varies among natural populations. We evaluated pollen viability responses to elevated temperature in two closely related taxa of yellow monkeyflowers across multiple geographic scales to determine whether heat sensitivity varies among species, between populations and across elevations.

**Methods:** We conducted three common garden growth chamber experiments. First, we compared pollen viability responses to heat stress between *Mimulus guttatus* and members of the *M. tilingii* species complex. Second, we tested whether broad elevational categories explained variation in heat responses. Third, we evaluated heat sensitivity among nine eastern Sierra Nevada populations of *M. guttatus* spanning an elevational gradient. Pollen viability was assessed following exposure to control and heat treatments.

**Key results:** Heat exposure reduced pollen viability in all experiments. In the species comparison experiment, pollen viability decreased more strongly in the high-elevation restricted *M. tilingii* species complex than the widespread *M. guttatus*. We found substantial population-level variation in pollen viability within *M. guttatus* but not members of the *M. tilingii* complex. Broad elevational categories did not explain this variation in pollen viability responses. However, within eastern Sierra Nevada populations, reductions in pollen viability became stronger with increasing source elevation. Several populations deviated from this trend, indicating reproductive responses to heat cannot be explained by elevation alone.

**Conclusions:** Pollen heat sensitivity varies substantially among species and populations of yellow monkeyflowers. Elevation alone is insufficient to predict pollen viability responses to elevated temperatures, suggesting that local adaptation or genetic variation may contribute to reproductive heat tolerance. Some populations of *M. guttatus* may contain greater capacity to maintain reproductive function under future warming, highlighting the importance of considering intraspecific variation when predicting plant responses to climate change.

## INTRODUCTION

Climate change is causing temperatures to rise globally and simultaneously increasing the intensity, duration, and frequency of heatwaves (Stillman, 2019; Perkins-Kirkpatrick and Lewis, 2020; IPCC, 2021; Seneviratne et al., 2021; Domeisen et al., 2023). While extreme temperatures associated with heatwaves are detrimental across life history stages (Jagadish et al., 2016), even moderately elevated temperatures disproportionally affect plant reproduction (Hedhly et al., 2009; Resentini et al., 2023; Lohani et al., 2025). Reproductive sensitivity to heat poses a major challenge for wild plant populations, as warming causes a significant decrease in fecundity (Zi et al., 2023) and can hinder a population’s ability to adapt or track shifting climatic conditions (Cinto Mejia and Wetzel, 2023; Tushabe et al., 2023). Despite this acute reproductive sensitivity, few studies have characterized the effects of heat on reproductive development in natural plant systems (Denney et al., 2026).

All stages of reproductive development may be affected by heat stress; however, the most sensitive stage for plants is pollen development (Zinn et al., 2010; Tushabe and Rosbakh, 2025; Arnold et al., 2026). Pollen is likely more sensitive because it is smaller and more exposed to the external environment than ovules (Chaturvedi et al., 2021). Heat stress can affect pollen formation during microsporogenesis by disrupting meiosis and chromosomal segregation (Bomblies et al., 2015; Lohani et al., 2025), by altering the nutritive tapetal development required for proper pollen formation (Chaturvedi et al., 2021; Resentini et al., 2023), or interrupting the mitotic stages required for fully developed pollen (Lohani et al., 2025). Additionally, heat exposure can lead to poor anther dehiscence (Zhang et al., 2021) or loss of mature pollen functionality (Chaturvedi et al., 2021; Lohani et al., 2025). While agricultural systems have been the focus of recent research on heat and pollen viability, we know far less about the impacts of heat stress on pollen viability in natural systems (Tushabe et al., 2023; Rosenberger et al., 2024).

Natural populations may vary substantially in their capacity to maintain reproductive function under elevated temperatures (Walsh et al., 2019). Local adaptation to thermal environments could generate population-level differences in pollen performance and enable populations from warmer environments to maintain pollen viability under heat stress (Jackwerth et al., 2024). Although relatively few studies have investigated this question in natural systems, research on wild relatives of crop species have demonstrated that there is often substantial genetic variation for reproductive responses to heat within species. For example, significant variation in pollen traits has been documented among populations of wild diploid potato (Nicolao et al., 2025), wild tomato relatives (Driedonks et al., 2018), and wild quinoa (Xu et al., 2025). These findings suggest that standing genetic variation for heat tolerance may be common in wild plant populations, but the extent of such variation remains poorly characterized.

Monkeyflowers provide an ideal system for examining variation in reproductive responses to heat stress. The common yellow monkeyflower (*Mimulus guttatus* syn. *Erythranthe guttata*; Phrymaceae) occurs across a broad range of environmental conditions in western North America, from sea level to subalpine habitats (Vickery, 1978). Populations of *M. guttatus* are locally adapted to their native environments (Lowry and Willis, 2010; DeMarche et al., 2016), making this species well suited for evaluating whether reproductive responses to heat vary among populations. In contrast, members of the *Mimulus tilingii* (syn. *Erythranthe tilingii*) species complex are closely related to *M. guttatus* but are restricted to high-elevation environments that generally experience cooler growing season temperatures. Together, these species provide an opportunity to test whether reproductive heat tolerance varies both among populations within species and between taxa that differ in their geographic and climatic distributions.

Here, we used three common-garden growth chamber experiments to examine how elevated temperatures affect pollen viability across species and populations of monkeyflowers. Specifically, we asked whether pollen viability declines under heat stress, whether high-elevation members of the *M.* tilingii complex are more sensitive to elevated temperatures than *M. guttatus*, and whether populations originating from warmer environments maintain greater pollen viability under heat stress than populations from cooler environments. We predicted that 1) exposure to heat would reduce pollen viability, 2) members of the *M. tilingii* complex would be more sensitive to heat than *M. guttatus* and 3) populations originating from lower elevation environments would maintain higher pollen viability under heat stress than populations from cooler, high-elevation environments.

## METHODS

### Study systems

The common yellow monkeyflower (*Mimulus guttatus* syn. *Erythranthe guttata*; Phrymaceae) is widespread throughout western North America where its range spans a diverse set of habitats across elevational gradients, ranging from sea level to subalpine meadows (Vickery, 1978). Across the diverse habitats that occur over its range, *M. guttatus* varies in life history strategies and mating systems. For example, outcrossing perennial populations are found near riversides and ocean bluffs while primarily selfing inland annuals are associated with ephemeral water availability (Lowry et al., 2008; Wu et al., 2008; Lowry and Willis, 2010; Friedman and Willis, 2013; Kooyers et al., 2025). Populations of *M. guttatus* are locally adapted (Lowry and Willis, 2010; DeMarche et al., 2016) and contain exceptionally high levels of genetic diversity (Puzey et al., 2017; Lovell et al., 2025), which can facilitate adaptation to changing climates (Kooyers et al., 2025).

The *Mimulus tilingii* (syn. *Erythranthe tilingii*) species complex is a group of high-elevation restricted yellow mountain monkeyflowers closely related to *M. guttatus*. Recent taxonomic revisions divided *M. tilingii* into three allopatric species (*M. caespitosa, M. minor,* and *M. tilingii*; Nesom, 2014) which are genetically distinct from one another (Sandstedt et al., 2021). Of these species, *M. tilingii* is widespread throughout western North America, while *M. caespitosa* (syn. *Erythranthe caespitosa*) grows in the Cascades and Olympic mountains, and *M. minor* (syn. *Erythranthe minor*) is primarily found in the Rocky Mountains of Colorado (Nesom, 2012). Members of the *M. tilingii* complex grow along alpine streams and can co-occur with populations of *M. guttatus*. Despite sympatry, hybrid seed lethality is a strong postzygotic barrier between *M. guttatus* and members of the *M. tilingii* complex, which may prevent hybridization between the two taxa (Garner et al., 2016). Further, while *M. tilingii* species are primarily outcrossing, recent evidence indicates some populations may have increased selfing rates (Sandstedt et al., 2021).

### Growth chamber experiments

To examine the effects of heat on pollen viability, we performed three separate experiments in a pair of growth chambers (BioChambers FXC-19/Spectral LED; Manitoba, Canada) outfitted with Phillips GreenPower LED modules. The first experiment used populations of the widespread *M. guttatus* and alpine restricted *M. tilingii* species complex to test whether high-elevation restricted species will have decreased pollen viability under elevated temperatures compared to their widespread congeners. This species comparison experiment suggested low elevation populations may contain genetic variation to buffer against heat stress. As a result, we conducted a second experiment to evaluate whether elevation of origin in *M. guttatus* plays a role in pollen viability in these populations (Fig. 1). Finally, we conducted a third growth chamber experiment, with increased population sampling in the southern populations of *M. guttatus*, to assess differences in pollen viability along an elevational gradient in the eastern Sierra Nevada Mountains (Fig. 2).

**Figure 1.**
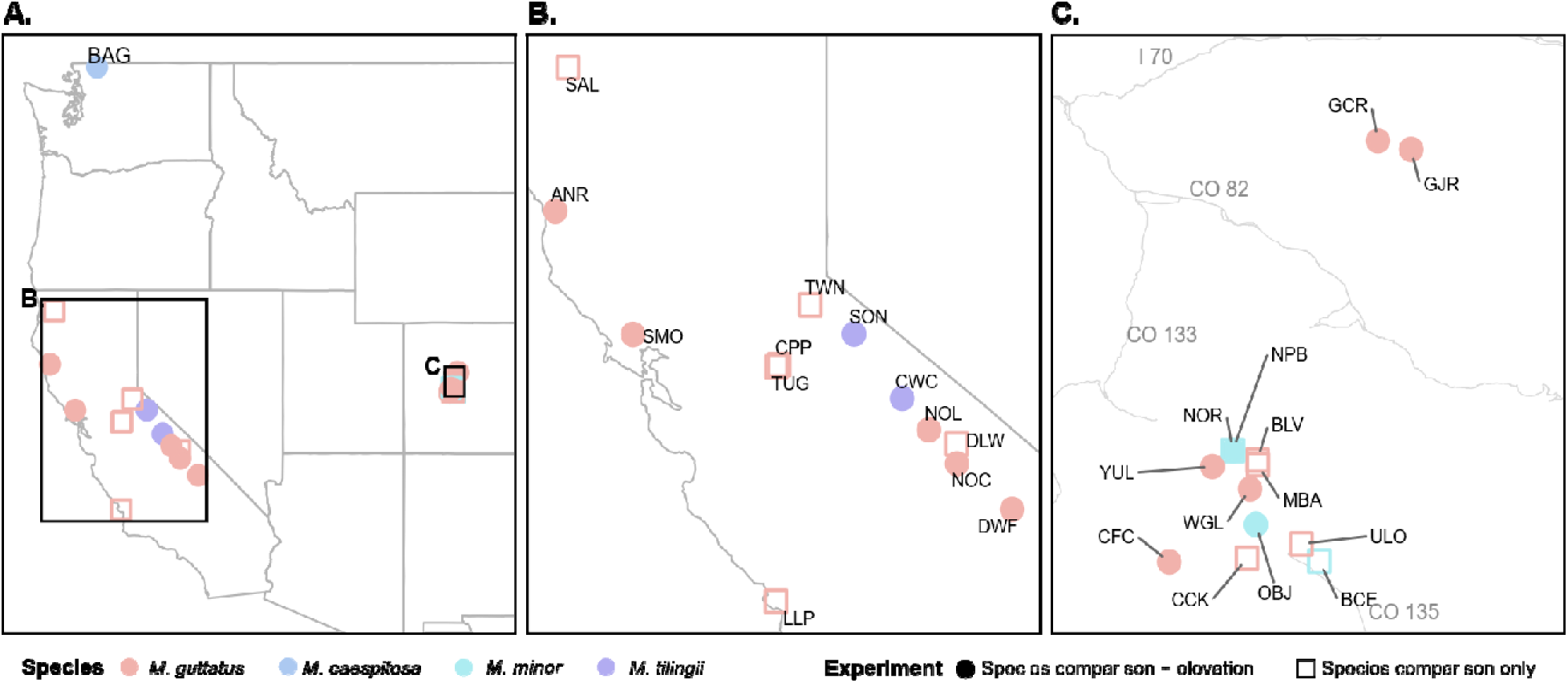
Map of *Mimulus guttatus* and *M. tilingii* species complex populations used in the species comparison and elevation experiments. Point color denotes species. Circles indicate populations used in both experiments, whereas open squares indicate populations used only in the species comparison experiment. Population abbreviations are labeled with three-letter codes. **A.** All populations included in the species comparison and elevation experiments across the western United States with inset boxes highlighting panels B and C. **B.** Populations from California. Gray lines indicate geographic boundaries. **C.** Populations from Colorado. Gray lines indicate local roadways. I= Interstate, CO = Colorado state highway.

**Figure 2.**
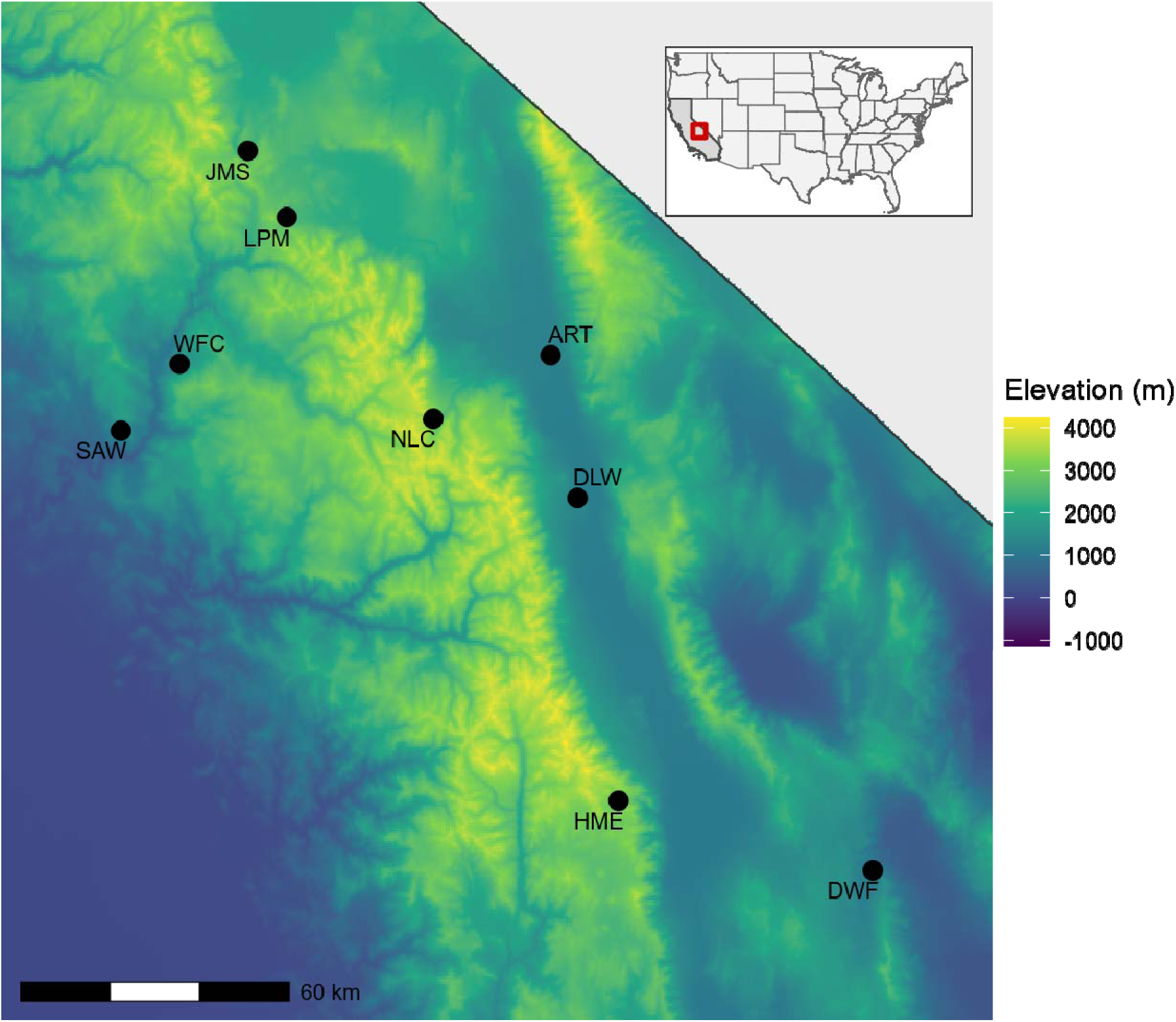
Map of *Mimulus guttatus* populations used in the eastern Sierra Nevada experiment. Black dots indicate population location. Population abbreviations are shown as three-letter codes. Elevation is shown as a color gradient, ranging from dark blue at lower elevations to yellow at higher elevations. The red bounding box in the inset map indicates the location of the study populations in California.

#### Species comparison experiment

We conducted a growth chamber common garden experiment to assess the hypothesis that members of the high-elevation restricted *M. tilingii* species complex have lower pollen viability when exposed to heat than their widespread congener, *M. guttatus*. We used a total of 27 populations of both species (n=20 *M. guttatus*; n=7 *M. tilingii* complex), including populations in relative sympatry, when available (Fig. 1).

From each population, we selected ten individuals, with five individuals randomly assigned to each treatment (n=27 populations x 5 individuals x 2 treatments = 270). All plants were germinated and grown in control conditions (16 h 22°C : 8 h 18°C, day : night) to minimize exposure to heat until floral buds had initiated. Across the range of *M. guttatus*, some southern populations can regularly experience air temperatures of 36°C during their reproductive phase. Therefore, we used 16 hour days with 36°C and 8 hour nights at 26°C as our heat treatment. As floral buds developed on mature plants, individual plants were moved to their preassigned, randomly chosen treatment and experimental flat where they were well-watered and monitored for flowering. From each plant, flowers were collected for pollen viability analysis (outlined below).

#### Elevation experiment

The results of the first experiment indicated there may be population-specific responses within *M. guttatus*, which could relate to source elevation. Therefore, we conducted our second growth chamber experiment to assess whether pollen viability differences were associated with elevation-of-origin in the widespread *M. guttatus.* Additionally, we asked whether high elevation populations of *M. guttatus* would have higher pollen viability than the high-elevation restricted *M. tilingii* species complex when exposed to heat during the reproductive phase.

For this elevation-of-origin experiment, we selected a subset of 15 populations from the species comparison experiment (Fig. 1) with five populations of *M. guttatus* from low elevations (mean elevation = 754 m), five populations from high elevations (mean elevation = 3054 m), and five populations from the *M. tilingii* complex (mean elevation = 2619 m). We used nine individuals per population per treatment for a total of 270 individuals (n = 9 x 15 populations x 2 treatments). Each plant was germinated and grown under control conditions (16 h 22°C : 8 h 18°C, day : night) until floral buds developed and then individuals randomly assigned to the heat treatment were grown in the heat chamber set to 30°C. This temperature reflects the mean maximum temperature across all lower elevation populations of *M. guttatus*. Ten days after the initial flowering date, we collected flower counts from each plant to determine whether increased temperature also affected total flower production.

#### Eastern Sierra Nevada experiment

The first two experiments relied upon populations from across the range of *M. guttatus*, which span latitudinal gradients and diverse mountain systems. To evaluate whether pollen viability might be locally adaptive at a regional scale, we conducted a third growth chamber experiment using *Mimulus guttatus* populations sourced from the eastern side of the southern Sierra Nevada mountain range (Fig. 2). This region of the Sierra Nevada mountains is one of the most topographically variable regions in the contiguous United States (Halofsky, 2021) where populations are primarily fed by prolonged snowmelt and remain wet during the growing season, which may temper the effects of heat exposure in these populations (Kimball et al., 2004).

For the third experiment, we selected nine populations from across this broad elevational gradient (24-3282m, mean = 1920.3m) to use in the experiment. For each population, we sowed 50-100 seeds in 2.5-inch square pots filled with a general propagation mix. Seeds were stratified for nine days at 4°C, after which they were monitored for germination in a growth chamber set for control conditions (16 h 22°C : 8 h 18°C, day : night; lights at 380mE). After germination, 20 seedlings were randomly chosen from each population and transplanted to individual 2.5-inch pots filled with sure-mix soil for use in the experiment (n= 9 populations x 10 individuals x 2 treatments = 180). Each pot was randomized into a flat and treatment. As floral buds developed on mature plants, individual plants were moved to an experimental flat either in the control conditions or the warming treatment (16 h 37°C : 8 h 26°C, day : night; lights at 380mE), where they were well-watered and monitored for flowering.

### Assessing pollen viability

Pollen staining using 0.1% aniline blue in lactophenol has been used as a proxy for viability in many plant species, including *Mimulus* (Sweigart et al., 2006; Yeamans et al., 2014; Seale, 2020; Stokes and Geitmann, 2025). Aniline blue stains well-developed cytoplasm dark blue while leaving underdeveloped pollen unstained or light blue (Fishman et al., 2013). To assess pollen viability, we collected four anthers from the second pair of flowers, as these pairs developed under their respective temperature treatment. Pollen was removed from each stamen by laying the anthers on a microscope slide and gently prodding the anther. We then added 15 μl of 0.1% lactophenol-aniline blue (Kearns and Inouye, 1993) directly onto the pollen, added a cover slip, and allowed the slide to dry for up to 24 hours. Each slide was imaged using an AmScope compound microscope with 40x magnification and a Nikon DS-Fi3 microscope camera with a field of view of 0.944 mm^2^. We randomly selected three non-overlapping sections of the slide for image capture.

We used the Cell Counter plug-in within FIJI (Schindelin et al., 2012) to manually count pollen for each captured image by considering dark blue pollen as viable and unstained or light gray grains as inviable. We calculated the total viable and inviable pollen counted across the three sampled images for each individual, as well as the proportion of viable pollen.

### Statistical analyses

We summarized our data and conducted statistical analyses using R v. 4.4.1 (RTeam, 2021). To test differences in pollen viability, we modeled the proportion of viable pollen out of all pollen counted using beta-binomial generalized linear mixed models in glmmTMB v. 1.1.11 (Brooks et al., 2017). Fixed effects were evaluated using Type III Wald chi-square tests with the car package v. 3.0-13 (Fox & Weisberg, 2019). Model fit was assessed using simulation-based residual diagnostics in DHARMa v. 0.4.7 (Hartig, 2022).

In the species comparison experiment, fixed effects included species (*M. guttatus* vs. *M. tilingii* complex), treatment, and their interaction, and population was included as a random intercept with a random slope for treatment to allow responses to vary among populations.

In the elevation-of-origin experiment, fixed effects included elevation type (low-elevation *M. guttatus*, high-elevation *M. guttatus*, and *M. tilingii* species complex), treatment, and their interaction, and population was included as a random intercept with a random slope for treatment. With this dataset, we also accounted for heteroscedasticity by modeling the dispersion parameter as a function of treatment.

In the eastern Sierra Nevada experiment, fixed effects included scaled source elevation (s_elev; z-scored to mean 0 and SD 1), treatment, and their interaction. We also accounted for variation within treatments with treatment blocks as a random effect. Although statistical models were fit using scaled elevation to improve numerical stability, we display our figures with raw elevation values for biological interpretability.

To evaluate reproductive performance in the elevation-of-origin experiment, we also analyzed flower production after ten days of flowering under the treatment conditions. Flower number was modeled using a negative binomial generalized linear mixed model in glmmTMB v. 1.1.11 (Brooks et al., 2017), with elevation category (low-elevation *M. guttatus*, high-elevation *M. guttatus*, or *M. tilingii* complex), treatment, and their interaction as fixed effects. Population and replicates were included as random intercepts to account for non-independence among individuals within populations.

To conduct population contrasts, we conducted post hoc inference by estimating marginal means and pairwise contrasts with emmeans v. 1.11.0 (Lenth, 2022). Treatment effects were summarized as rate ratios by exponentiating log-scale contrasts, and p-values for within-population contrasts were adjusted using the Holm method.

## RESULTS

### Species comparison experiment

Heat strongly reduced pollen viability in both *M. guttatus* and the *M. tilingii* species complex (Fig. 3). Estimated marginal means indicated that pollen viability declined from 87.3% to 17.1% in *M. guttatus* and from 90.0% to 4.1% in the *M. tilingii* species complex, corresponding to a decline of approximately 80% and 96%, respectively. The greater reduction in pollen viability in the *M. tilingii* complex was supported by a significant species-by-treatment interaction (χ^2^=4.50, df = 1, *P=*0.034) in the beta-binomial mixed model (Fig. 3, Table 1). Post hoc population-specific contrasts showed that heat significantly reduced pollen viability in most populations, although the magnitude of this effect varied among *M. guttatus* populations (Fig. S1).

**Figure 3.**
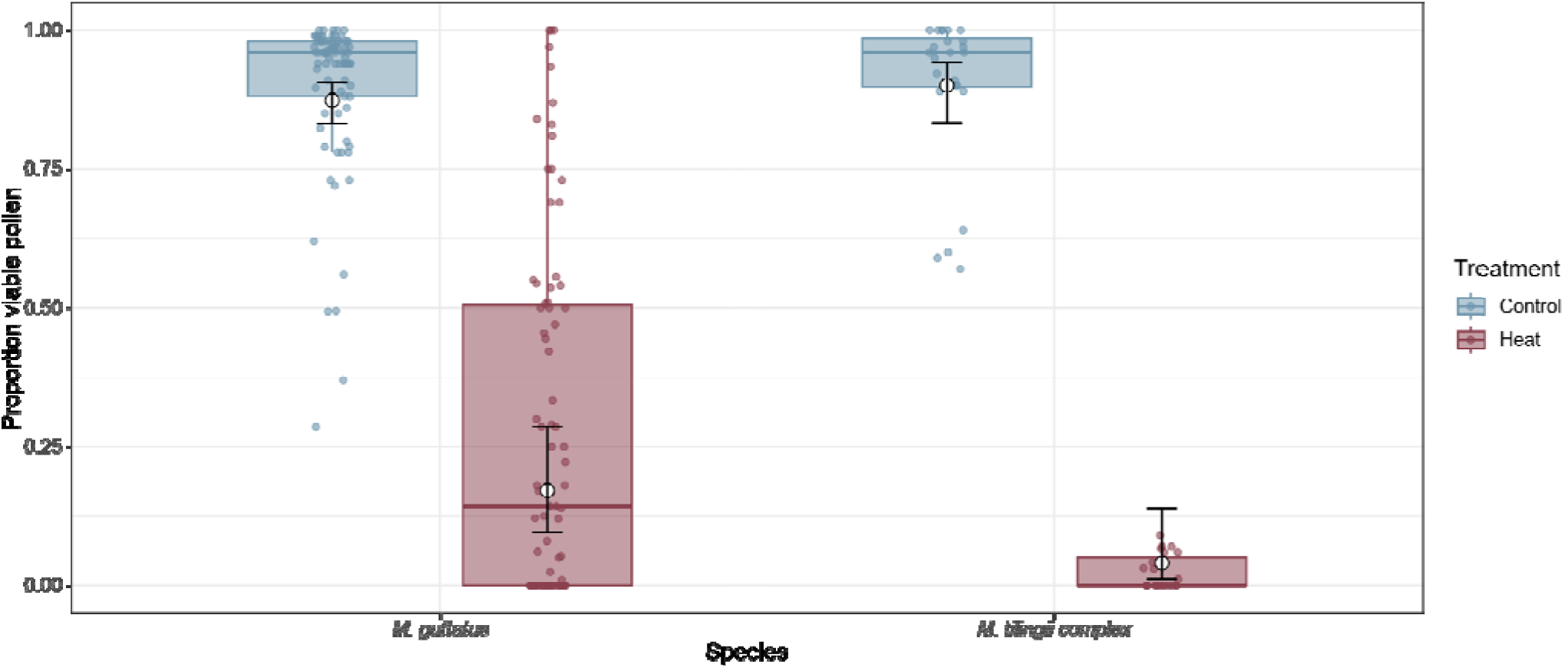
Heat substantially reduced pollen viability in both *Mimulus guttatus* and the *M. tilingii* species complex at 36°C. Raw data points are represented by points and summarized by boxplots. White dots and error bars indicate the model-estimated marginal means and 95% confidence intervals from the beta binomial generalized linear mixed model.

**Table 1.** ANOVA tables representing proportion viable pollen for each experiment. Significant p-values are in bold.

| Species Comparison Experiment |  |  |  | Elevation Experiment |  |  |  | Eastern Sierra Nevada Experiment |  |  |  |
| --- | --- | --- | --- | --- | --- | --- | --- | --- | --- | --- | --- |
| Proportion viable pollen |  |  |  | Proportion viable pollen |  |  |  | Proportion viable pollen |  |  |  |
| Predictor | $\chi^2$ | df | *P* | Predictor | $\chi^2$ | df | *P* | Predictor | $\chi^2$ | df | *P* |
| Species | 0.69 | 1 | 0.41 | Elevation category | 0.94 | 2 | 0.62 | Source elevation | 0.22 | 1 | 0.64 |
| Temperature | 71.41 | 1 | **<0.0001** | Temperature | 34.65 | 1 | **<0.0001** | Temperature | 59.43 | 1 | **<0.0001** |
| Species x Temperature | 4.5 | 1 | **0.034** | Elevation category x Temperature | 0.97 | 2 | 0.61 | Source elevation x Temperature | 4.17 | 1 | **0.041** |

### Elevation experiment

Heat significantly reduced pollen viability at 30°C (Fig. 4A; Table 1; χ^2^=34.65, df = 1, *P*<0.0001). Pollen viability in the low-elevation populations of *M. guttatus* declined from 84.7% to 60.4%, the high-elevation populations decreased from 88.7% to 60.2%, and the members of the *M. tilingii* species complex were reduced from 91.2% to 66.7%. However, the magnitude of these reductions was not significantly related to elevation (χ^2^=0.94, df = 2, *P=*0.62), nor did we recover a significant interaction between temperature and elevation (χ^2^ = 0.97, df = 2, *P*=0.61). This indicates the lower heat treatment affected pollen viability similarly across all populations (Fig. 4A, Table 1).

**Figure 4.**
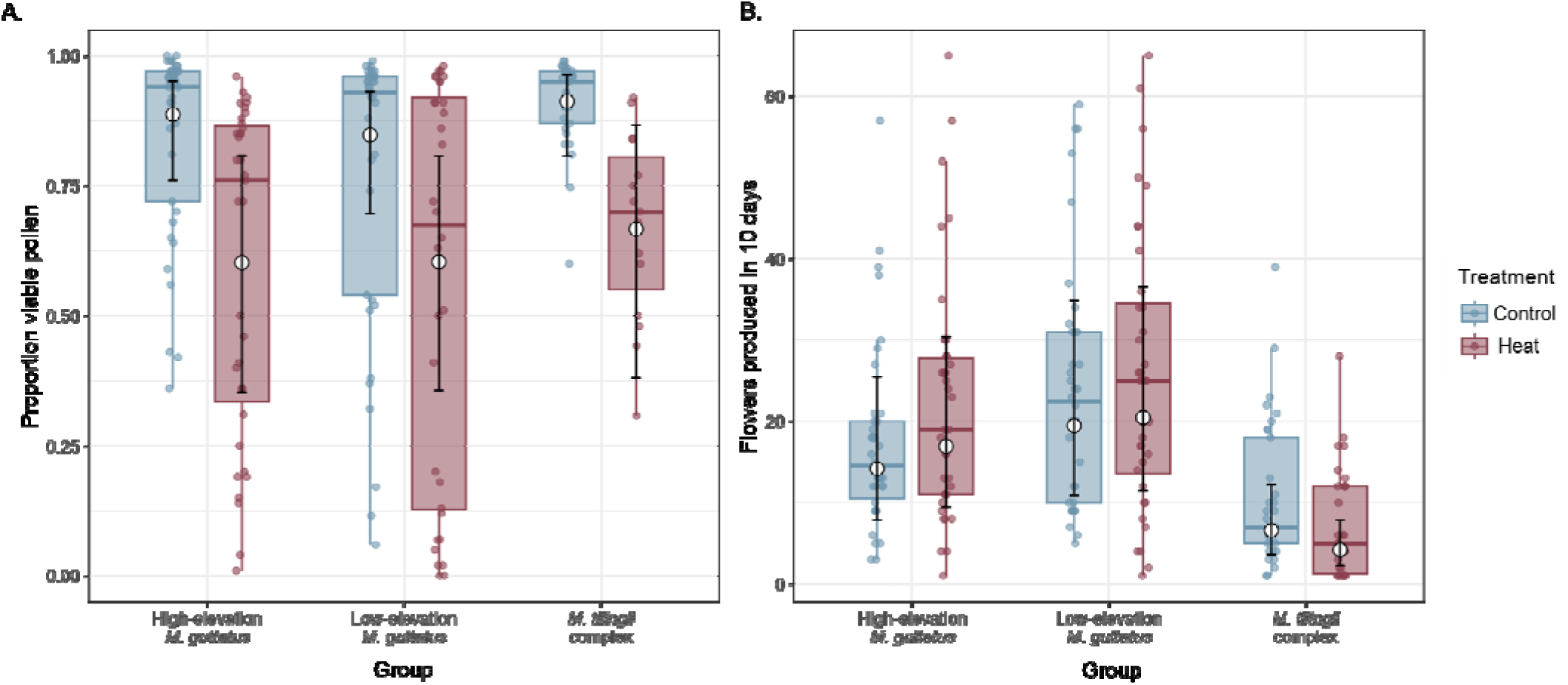
Heat effect on pollen viability and flower number across elevational categories at 30°C. A. Pollen viability was significantly affected by temperature but did not vary among elevational categories. B. Flower number after 10 days was unchanged by heat in high- and low-elevation *M. guttatus* but significantly decreased in the *M. tilingii* complex. Filled points show raw data and boxplots summarize distributions. White dots and error bars indicate the model-estimated marginal means and 95% confidence intervals.

Flower production responded differently to heat among the three groups (Fig 4B; Table 2). Neither the low nor high elevation *M. guttatus* groups had a significant change in flower number; however, members of the *M. tilingii* complex decreased floral production by 36.5% under heat treatment. These differences were reflected in a significant treatment by group interaction (χ^2^=12.96, df=2, *P*=0.0015). Post hoc comparisons confirmed that heat did not significantly alter flower number in the high- or low-elevation *M. guttatus* populations, whereas members of the *M. tilingii* complex produced significantly fewer flowers under heat than under control (Control/Heat = 1.57, P=0.0014).

**Table 2.** ANOVA table indicating the number of flowers 10 days after heat treatment in the elevation experiment. Significant p-values are in bold.

| Flower Number 10 days post treatment |  |  |  |
| --- | --- | --- | --- |
| Predictor | $\chi^2$ | df | *P* |
| Elevation category | 6.58 | 2 | **0.037** |
| Treatment | 2.63 | 1 | 0.11 |
| Elevation category<br>x Treatment | 12.96 | 2 | **0.0015** |

**Table 3.**
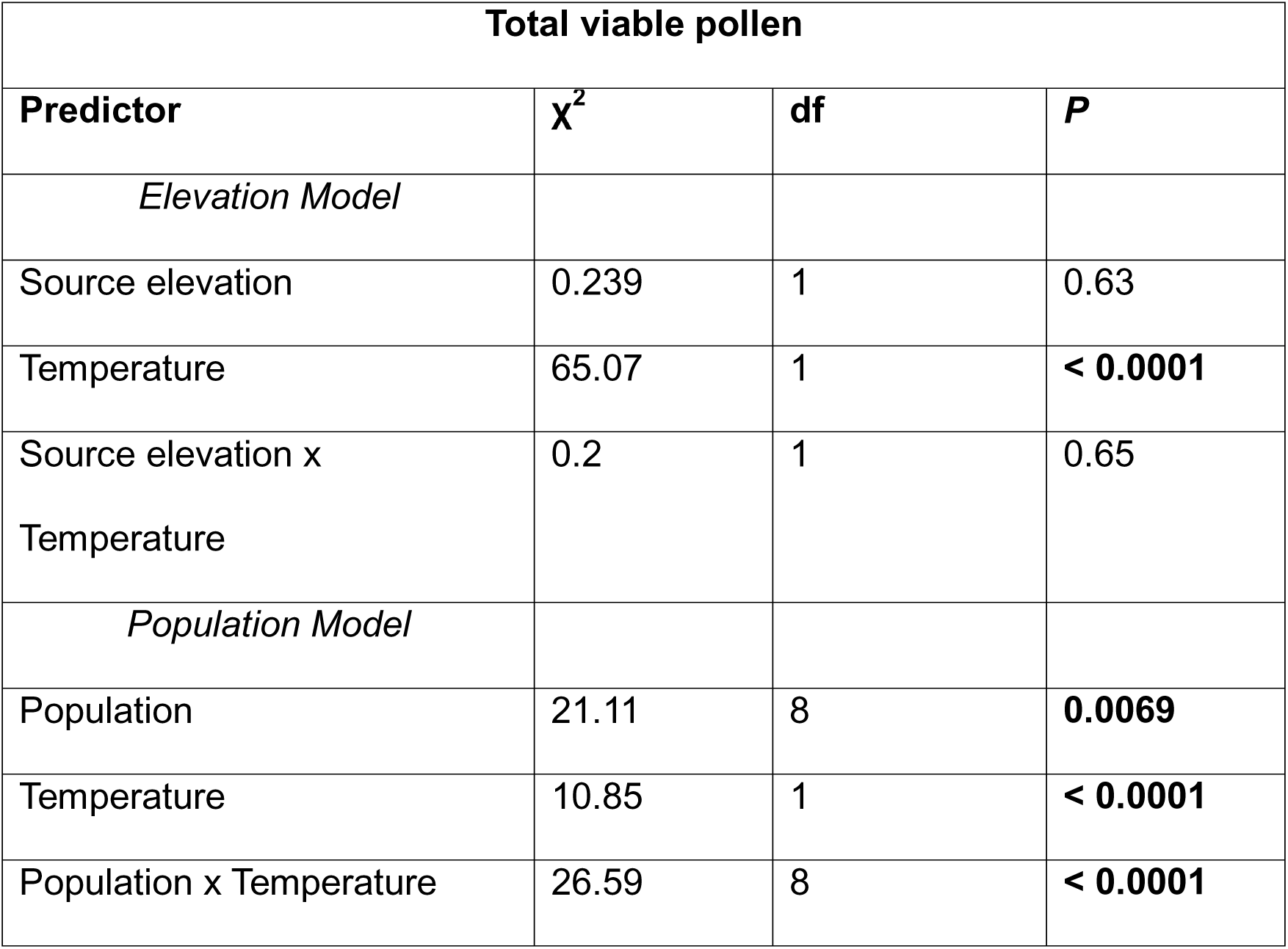
ANOVA tables for total viable pollen counts. The elevation model used source elevation as a factor whereas the population model included population codes without source elevation. Significant p-values are bolded.

| Total viable pollen |  |  |  |
| --- | --- | --- | --- |
| Predictor | $\chi^2$ | df | *P* |
| *Elevation Model* |  |  |  |
| Source elevation | 0.239 | 1 | 0.63 |
| Temperature | 65.07 | 1 | **< 0.0001** |
| Source elevation x<br>Temperature | 0.2 | 1 | 0.65 |
| *Population Model* |  |  |  |
| Population | 21.11 | 8 | **0.0069** |
| Temperature | 10.85 | 1 | **< 0.0001** |
| Population x Temperature | 26.59 | 8 | **< 0.0001** |

### Eastern Sierra Nevada experiment

As with the other experiments, pollen viability was strongly reduced by heat treatment (Table 1; χ^2^ = 59.43, df = 1, *P* < 0.0001). Predicted pollen viability declined by approximately 29% in low-elevation populations and 42% in high-elevation populations under heat treatment. We found a significant interaction between treatment and source elevation (χ^2^ = 4.17, df = 1, *P =* 0.041, β = −0.307), indicating that the heat-associated reductions in pollen viability became stronger with increasing elevation (Fig. 5).

**Figure 5.**
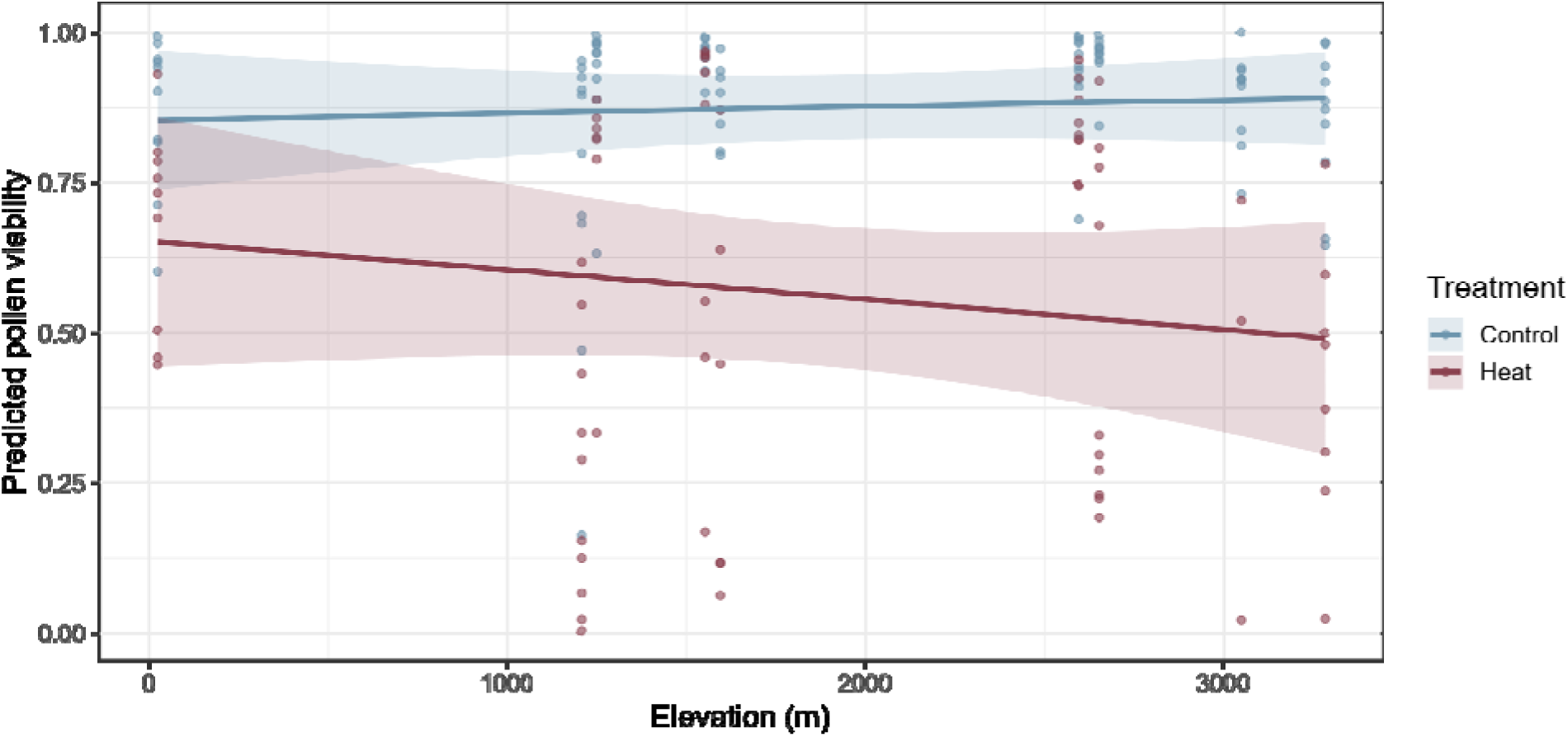
Predicted pollen viability by elevation in eastern Sierra Nevada populations of *Mimulus guttatus* decreased with elevation. Pollen viability declined with elevation and was lower under heat than control. Lines show model-predicted values with shaded regions indicating 95% confidence intervals. Points indicate raw data observations.

We also found that total pollen production was strongly reduced by heat in the eastern Sierra Nevada populations of *M. guttatus* (χ^2^ = 10.85, df = 1, *P* < 0.001) although the magnitude of reduction varied significantly among populations (χ^2^ = 26.59, df = 8, *P* < 0.0001). Within populations, heat effects were summarized using estimated marginal means as rate ratios (Heat / Control; Fig. 6; Fig. S2). Rate ratios ranged from 0.089 in ART (∼91% reduction) to 0.714 in DLW (∼ 29% reduction). After correction across the nine within-population contrasts, heat significantly reduced total pollen in populations DWF, SAW, ART, JMS, HME and NLC, but had no significant decrease in DLW, WFC, and LPM.

**Figure 6.**
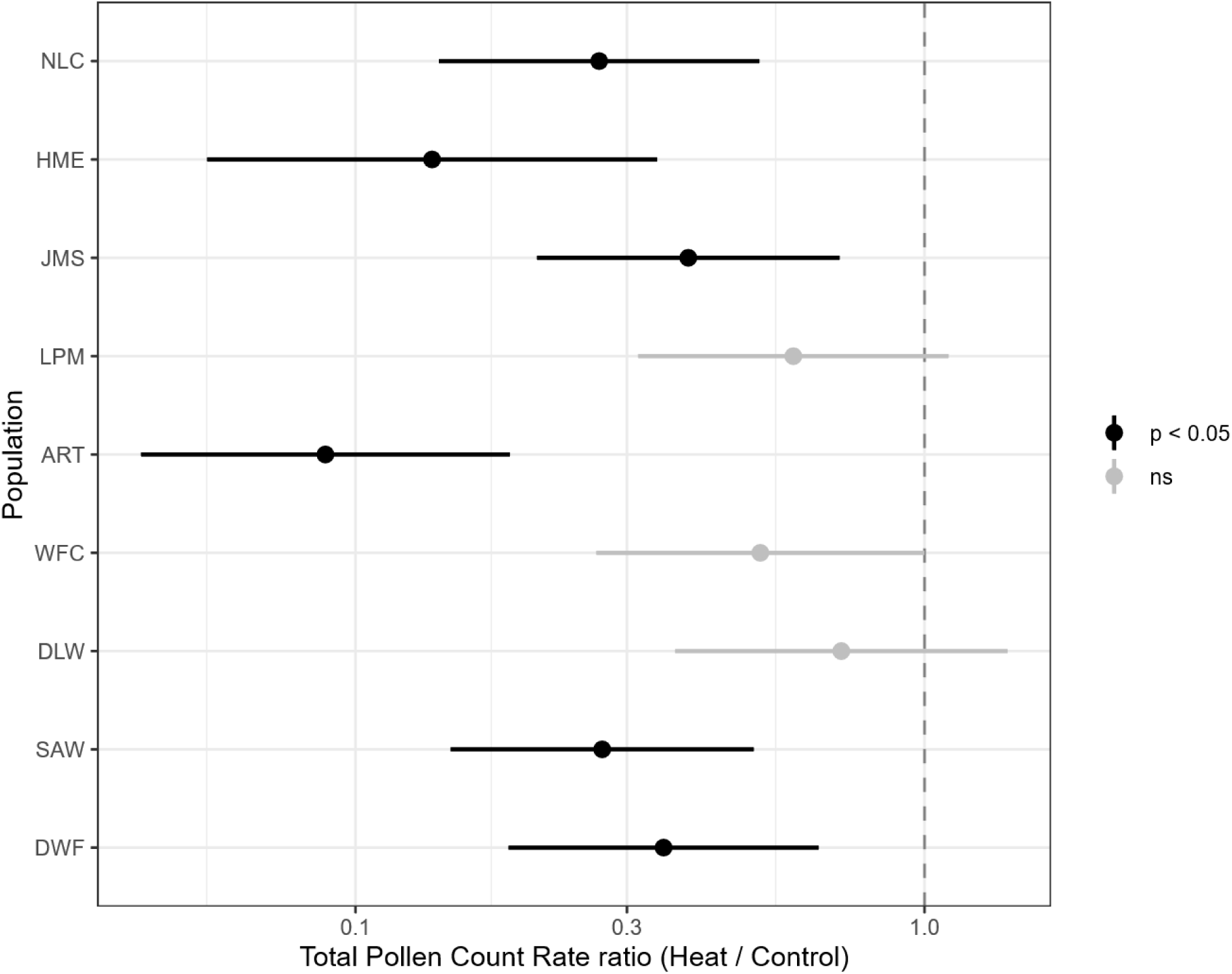
Population-specific effects of heat exposure on sampled pollen counts in eastern Sierra Nevada populations of *Mimulus guttatus*. Points show estimated heat to control rate ratios for the total number of pollen grains counted across standardized images of each population. Horizontal bars indicate 95% confidence intervals. Black points denote significant treatment effects within populations and gray points indicate non-significant effects. Populations are arranged along the y-axis from lowest elevation (DWF) to highest elevation (NLC).

## DISCUSSION

We performed a series of common garden growth chamber experiments to evaluate how elevated temperatures affect pollen viability in *Mimulus guttatus* and members of the *M. tilingii* species complex across multiple geographic scales. Heat exposure generally reduced pollen viability, but the magnitude of the reduction varied among species and source populations. In our species comparison experiment, we detected substantial population-level variation in *M. guttatus* but not in the *M. tilingii* complex (Fig. 3). Contrary to our predictions, variation in *M. guttatus* was not explained by broad elevational categories (Fig. 4). However, in the eastern Sierra Nevada experiment, reductions in pollen viability under heat became stronger with increasing source elevation (Fig. 5). Several populations deviated from this overall pattern, indicating that reproductive responses to heat cannot be explained by source elevation alone (Figs. 5, 6). Collectively, these findings suggest populations differ in their ability to maintain reproductive function under elevated temperatures and are consistent with the presence of genetic variation for heat sensitivity within *M.* guttatus.

### Population-level variation in heat sensitivity

Among *M. guttatus* populations, responses to elevated temperature were highly variable. We predicted low-elevation populations would be less susceptible to heat than high-elevation populations. Contrary to our prediction, broad elevational categories did not explain the high variability we found across populations, as low- and high-elevation populations performed similarly in the elevation-of-origin experiment. However, the eastern Sierra Nevada experiment revealed a significant relationship between source elevation and pollen viability under heat stress, with higher elevation populations generally exhibiting larger reductions in viability. Nonetheless, several populations deviated from this expectation in surprising ways. For example, the low elevation SAW population (1208 m) experienced one of the largest declines in pollen viability under heat stress while the low-elevation ART population (1594 m) showed the largest decline in total pollen production. Additionally, the high-elevation LPM population (2594 m) maintained relatively high pollen viability and an insignificant decrease in total pollen production under heat stress, despite its high-elevation origins. These contrasting responses suggest that some populations may harbor loci that contribute to maintaining reproductive function under elevated temperatures. As such, gene flow among populations could facilitate the spread of heat-tolerant alleles and enhance adaptive responses to warming environments. Indeed, demographic experiments indicate low-elevation populations of *Mimulus guttatus* may have upslope migration potential and be favored under selection in high-elevation locations (DeMarche et al., 2016; DeMarche et al., 2020). However, we know little about the migratory potential of the high-elevation populations. Additionally, recent work has identified reproductive isolation among some *M. guttatus* lineages (Lovell et al., 2025; Frayer et al., 2026), which may constrain introgression from these populations. Future studies that incorporate genomic data and reproductive performance under heat stress can help evaluate the extent to which reproductive heat tolerance can spread among populations through gene flow.

In contrast to *M. guttatus,* members of the *M. tilingii* species complex exhibited strong reductions in pollen viability, little evidence of population-specific responses and a significant decrease in flower numbers in response to elevated temperatures. Because members of the *M. tilingii* complex are predominantly outcrossing, reductions in both flower production and pollen viability may decrease opportunities for successful fertilization under elevated temperatures.

### Implications for adaptation under climate change

Population-level differences in pollen viability under heat stress may have important consequences for reproductive success under future warming. Pollen development is widely recognized as one of the most heat-sensitive stages of the plant life cycle and reductions in pollen viability can translate to reduced fertilization success and lower seed production (Chaturvedi et al., 2021; Lohani et al., 2025; Arnold et al., 2026). Most studies examining these relationships have focused on crop systems, where reproductive failure is a major driver of yield loss (Tushabe and Rosbakh, 2025). However, recent studies using wild plant species have revealed substantial variation in pollen responses to heat stress. For example, pollen performance differs between lowland and highland populations of *Lotus corniculatus* (Fabaceae), suggesting local adaptation of reproductive traits to thermal environments in this system (Jackwerth et al., 2024). Hsu and Kim (2025) found that pollen performance was associated with elevation of origin only in *Pinus contorta* but not in two other pine species, indicating that elevational patterns in reproductive traits are not necessarily consistent across taxa. In contrast, Heiling and Koski (2024) reported unexpectedly high pollen performance in some high-elevation populations of the forb, *Argentina anserina* (Rosaceae). Our results extend these findings by showing substantial variation in pollen viability responses to heat stress among populations of *Mimulus guttatus*. Collectively, these results indicate that pollen performance in natural systems may respond in diverse and unexpected ways to temperature stress.

The contrasting responses of *M. guttatus* and the *M. tilingii* complex have important implications for adaptation under future climate change. The substantial variation observed among *M. guttatus* populations suggests that some populations may harbor alleles that contribute to the maintenance of reproductive function under elevated temperatures. If these differences have a genetic basis, gene flow among populations could facilitate the spread of heat-tolerant alleles and enhance adaptive responses to warming environments. However, the movement of adaptive variation may be constrained by population structure and reproductive barriers. Recent work has identified substantial genetic differentiation and reproductive isolation among some *M. guttatus* lineages (Frayer et al., 2026), suggesting that adaptive variants may not move freely throughout the species. Opportunities for adaptive introgression may be even more limited in the *M. tilingii* complex. Although *M. guttatus* and members of the *M. tilingii* complex can occur in sympatry, strong postzygotic reproductive barriers substantially reduce gene exchange between these taxa (Sandstedt et al., 2021). Consequently, adaptation to future warming may depend largely on standing genetic variation already present within populations.

Our study focused on male reproductive performance as measured by pollen viability. Although pollen viability is widely used as a proxy for male fertility, successful reproduction also depends on pollen germination, pollen tube growth, ovule development, and seed production. Future studies linking pollen viability to realized seed production and fitness under warming temperatures will be necessary to determine whether the patterns observed here translate into meaningful demographic consequences. In addition, integrating reproductive heat responses with flowering phenology may improve predictions of how populations will experience and respond to future warming expected under climate change.

## CONCLUSIONS

Heat stress consistently reduced pollen viability across three experiments using *Mimulus guttatus* and members of the *M. tilingii* species complex, demonstrating the sensitivity of pollen development to elevated temperatures. However, responses were highly variable among species and populations, indicating reproductive vulnerability to warming is not uniform. Population-level variation within *M. guttatus* suggests that some populations may possess greater capacity to maintain reproductive function under future warming, whereas members of the *M. tilingii* complex exhibited consistently negative reproductive responses to elevated temperatures. These findings highlight the importance of considering intraspecific variation in reproductive heat sensitivity when evaluating the impacts of climate change on natural plant populations.

## Author contributions

DAD and CNB designed and performed experiments and analyzed data. DBL conceived of the study. DAD wrote the original manuscript draft. All authors read and approved the final manuscript.

## Acknowledgements

Funding for this research was provided by Michigan State University through a Plant Resilience Institute postdoctoral fellowship to DAD and start-up funds to DBL. We thank Daniel Lowry, Damian Popovic, Madison Plunkert, and Sylvie Martin-Eberhardt for assistance with seed collections on the eastern side of the Sierra Nevada Mountain range. Carrie Wu and Andrea Sweigart graciously provided seeds of *M. tillingii* that made this research possible. Cody Keilen and Nick Deason provided growth chamber support. Emily Josephs, Magie Williams, Maya Wilson-Brown, and Andrew Fairclough provided valuable comments that helped to improve the manuscript.

## Data Availability

Data and R scripts used in this study are available on GitHub at: https://github.com/ddenney1/mimulus_pollen.

## Notes

### Competing Interest Statement

The authors have declared no competing interest.

## References

Arnold, P. A., T. J. Walker, E. V. Wishart, and L. K. Guja. 2026. Evaluating the vulnerability of critical early life stages in plants during heat extremes. Conserv Physiol 14: coag015.

Bomblies, K., J. D. Higgins, and L. Yant. 2015. Meiosis evolves: adaptation to external and internal environments. New Phytologist 208: 306–323 (en).

Chaturvedi, P., A. J. Wiese, A. Ghatak, L. Záveská Drábková, W. Weckwerth, and D. Honys. 2021. Heat stress response mechanisms in pollen development. New Phytologist 231: 571–585 (lb).

Cinto Mejia, E., and W. C. Wetzel. 2023. The ecological consequences of the timing of extreme climate events. Ecol Evol 13: e9661.

DeMarche, M. L., K. M. Kay, and A. L. Angert. 2016. The scale of local adaptation in Mimulus guttatus: comparing life history races, ecotypes, and populations. New Phytol 211: 345–356.

DeMarche, M. L., A. L. Angert, and K. M. Kay. 2020. Experimental migration upward in elevation is associated with strong selection on life history traits. Ecol Evol 10: 612–625.

Denney, D. A., A. L. Taylor, E. B. Josephs, J. H. Willis, and D. B. Lowry. 2026. The incredible vulnerability that reproduction poses for plant species in a warming world. New Phytol 10.1111/nph.71422 DOI.

Domeisen, D. I. V., E. A. B. Eltahir, E. M. Fischer, R. Knutti, S. E. Perkins-Kirkpatrick, C. Schär, S. I. Seneviratne, et al. 2023. Prediction and projection of heatwaves. Nature Reviews Earth & Environment 4: 36–50.

Driedonks, N., M. Wolters-Arts, H. Huber, G.-J. de Boer, W. Vriezen, C. Mariani, and I. Rieu. 2018. Exploring the natural variation for reproductive thermotolerance in wild tomato species. Euphytica 214.

Fishman, L., A. Stathos, P. M. Beardsley, C. F. Williams, and J. P. Hill. 2013. Chromosomal rearrangements and the genetics of reproductive barriers in mimulus (monkey flowers). Evolution 67: 2547–2560.

Frayer, M. E., H. K. Soliman, P. F. Schwarz, and J. M. Coughlan. 2026. Introgression and parental conflict shape repeated occurrences of postzygotic isolation in Mimulus. Current Biology 10.1016/j.cub.2026.02.022 DOI.

Friedman, J., and J. H. Willis. 2013. Major QTLs for critical photoperiod and vernalization underlie extensive variation in flowering in the Mimulus guttatus species complex. New Phytol 199: 571–583.

Garner, A. G., A. M. Kenney, L. Fishman, and A. L. Sweigart. 2016. Genetic loci with parent-of-origin effects cause hybrid seed lethality in crosses between Mimulus species. New Phytol 211: 319–331.

Halofsky, J. E. 2021. Climate change effects in the Sierra Nevada. USDA Forest Service PSW-GTR-272, P. S. R. Station, Albany, CA.

Hedhly, A., J. I. Hormaza, and M. Herrero. 2009. Global warming and sexual plant reproduction. Trends Plant Sci 14: 30–36.

Heiling, J. M., and M. H. Koski. 2024. Divergent gametic thermal performance and floral warming across an elevation gradient. Evolution 78: 665–678 (en).

Hsu, H. W., and S. H. Kim. 2025. Temperature dependence of pollen germination and tube growth in conifers relates to their distribution along an elevational gradient in Washington State, USA. Ann Bot 135: 277–292.

IPCC. 2021. Climate Change 2021: The Physical Science Basis. Contribution of Working Group I to the Sixth Assessment Report of the Intergovernmental Panel on Climate Change, Cambridge, United Kingdom and New York, NY, USA.

Jackwerth, K., P. Biella, and J. Klecka. 2024. Pollen thermotolerance of a widespread plant, Lotus corniculatus, in response to climate warming: possible local adaptation of populations from different elevations. PeerJ 12: e17148.

Jagadish, S. V., R. N. Bahuguna, M. Djanaguiraman, R. Gamuyao, P. V. Prasad, and P. Q. Craufurd. 2016. Implications of High Temperature and Elevated CO2 on Flowering Time in Plants. Front Plant Sci 7: 913.

Kearns, C., and D. Inouye. 1993. Techniques for Pollination Biologists. University Press of Colorado.

Kimball, S., P. Wilson, and J. Crowther. 2004. Local ecology and geographic ranges of plants in the Bishop Creek watershed of the eastern Sierra Nevada, California, USA. Journal of Biogeography 31: 1637–1657.

Kooyers, N. J., J. T. Anderson, A. L. Angert, M. L. Avolio, D. R. Campbell, M. Exposito-Alonso, T. E. Juenger, et al. 2025. Responses to climate change – insights and limitations from herbaceous plant model species. New Phytologist 248: 461–493.

Lohani, N., M. B. Singh, and P. L. Bhalla. 2025. Deciphering the vulnerability of pollen to heat stress for securing crop yields in a warming climate. Plant, Cell & Environment 48: 2549–2580 (en).

Lovell, J. T., R. Walstead, A. Lawrence, E. Stark-Dykema, M. C. Farnitano, A. Harder, T. Bruna, et al. 2025. Comparative Analyses of Four Reference Genomes Reveal Exceptional Diversity and Weak Linked Selection in the Yellow Monkeyflower (Mimulus guttatus) Complex. Mol Ecol Resour 25: e70012.

Lowry, D. B., and J. H. Willis. 2010. A widespread chromosomal inversion polymorphism contributes to a major life-history transition, local adaptation, and reproductive isolation. PLoS Biol 8.

Lowry, D. B., R. C. Rockwood, and J. H. Willis. 2008. Ecological reproductive isolation of coast and inland races of *Mimulus guttatus*. Evolution 62: 2196–2214.

Nesom, G. L. 2012. Taxonomy of Erythranthe sect. Simiola (Phrymaceae) in the USA and Mexico. Phytoneuron 40: 1–123.

Nesom, G. L. 2014. Updated classification and hypothetical phylogeny of Erythranthe sect. Simiola (Phrymaceae). Phytoneuron 81: 1–6.

Nicolao, R., I. Bashir, C. M. Castro, and G. Heiden. 2025. Evaluation of diploid wild potatoes pollen traits under heat stress. Potato Research 10.1007/s11540-025-09878-6 DOI(en).

Perkins-Kirkpatrick, S. E., and S. C. Lewis. 2020. Increasing trends in regional heatwaves. Nat Commun 11: 3357.

Puzey, J. R., J. H. Willis, and J. K. Kelly. 2017. Population structure and local selection yield high genomic variation in Mimulus guttatus. Mol Ecol 26: 519–535.

Resentini, F., G. Orozco-Arroyo, M. Cucinotta, and M. A. Mendes. 2023. The impact of heat stress in plant reproduction. Frontiers in Plant Science 14: 1271644 (en).

Rosenberger, N. M., J. A. Hemberger, and N. M. Williams. 2024. Heatwaves exacerbate pollen limitation through reductions in pollen production and pollen vigour. AoB Plants 16: plae045 (en).

Sandstedt, G. D., C. A. Wu, and A. L. Sweigart. 2021. Evolution of multiple postzygotic barriers between species of the Mimulus tilingii complex. Evolution 75: 600–613.

Schindelin, J., I. Arganda-Carreras, E. Frise, V. Kaynig, M. Longair, T. Pietzsch, S. Preibisch, et al. 2012. Fiji: an open-source platform for biological-image analysis. Nat Methods 9: 676–682.

Seale, M. 2020. Callose Deposition during Pollen Development. Plant Physiol 184: 564–565.

Seneviratne, S. I., X. Zhang, M. Adnan, W. Badi, C. Dereczynski, A. Di Luca, S. Ghosh, et al. 2021. Weather and Climate Extreme Events in a Changing Climate. *In* V. Masson-Delmotte, P. Zhai, A. Pirani, S.L. Connors, C. Péan, S. Berger, N. Caud, et al. [eds.], Climate Change 2021 – The Physical Science Basis, 1513–1766. Cambridge University Press, Cambridge, United Kingdom and New York, NY, USA.

Stillman, J. H. 2019. Heat Waves, the New Normal: Summertime Temperature Extremes Will Impact Animals, Ecosystems, and Human Communities. Physiology 34: 86–100.

Stokes, M., and A. Geitmann. 2025. Screening methods for thermotolerance in pollen. Ann Bot 135: 71–88.

Sweigart, A. L., L. Fishman, and J. H. Willis. 2006. A simple genetic incompatibility causes hybrid male sterility in mimulus. Genetics 172: 2465–2479.

Team, R.C. 2021. R: A Language and Environment for Statistical Computing. website: https://www.R-project.org/.

Tushabe, D., and S. Rosbakh. 2025. Patterns and drivers of pollen temperature tolerance. Plant, Cell and Environment 48: 1366–1379 (en).

Tushabe, D., F. Altmann, E. Koehler, S. Woods, and S. Rosbakh. 2023. Negative effects of high-temperature stress on gametophyte performance and their consequences for seed reproduction in wild plants. Environmental and Experimental Botany 216(English).

Vickery, R. K. 1978. Case Studies in the Evolution of Species Complexes in Mimulus. *In* M. K. Hecht, Steere, W.C., Wallace, B. [ed.], Evolutionary Biology, 405–507. Springer, Boston, MA.

Walsh, B. S., S. R. Parratt, A. A. Hoffmann, D. Atkinson, R. R. Snook, A. Bretman, and T. A. R. Price. 2019. The impact of climate change on fertility. Trends in Ecology & Evolution 34: 249–259 (English).

Wu, C. A., D. B. Lowry, A. M. Cooley, K. M. Wright, Y. W. Lee, and J. H. Willis. 2008. Mimulus is an emerging model system for the integration of ecological and genomic studies. Heredity (Edinb) 100: 220–230.

Xu, J., H. U. Farooq, M. Hashim, E. Rey, R. Curti, A. Morris, P. J. Maughan, et al. 2025. Wild relatives to improve heat tolerance of cultivated quinoa (Chenopodium quinoa): pollen viability and grain number. J Exp Bot 76: 5117–5128.

Yeamans, R. L., T. H. Roulston, and D. E. Carr. 2014. Pollen quality for pollinators tracks pollen quality for plants in Mimulus guttatus. Ecosphere 5: 1–8.

Zhang, Z., M. Hu, W. Xu, Y. Wang, K. Huang, C. Zhang, and J. Wen. 2021. Understanding the molecular mechanism of anther development under abiotic stresses. Plant Molecular Biology 105: 1–10 (de).

Zi, H., X. Jing, A. Liu, X. Fan, S.-C. Chen, H. Wang, and J.-S. He. 2023. Simulated climate warming decreases fruit number but increases seed mass. Global Change Biology 29: 841–855.

Zinn, K. E., M. Tunc-Ozdemir, and J. F. Harper. 2010. Temperature stress and plant sexual reproduction: uncovering the weakest links. J Exp Bot 61: 1959–1968.

