## Supplement for "Heat stress reveals divergent pollen viability responses across species and populations of monkeyflower"

Supplemental tables and figures

Table S1. Populations used in growth chamber experiments. Each population is identified with a three-letter code. Latitude and longitude are provided using the WGS84 datum.

| **Population** | **Species** | **Latitude** | **Longitude** | **Elevation (m)** | **A** | **B** | **C** |
| --- | --- | --- | --- | --- | --- | --- | --- |
| BAG | *M. caespitosa* | 48.87 | -121.69 | 1,279 | X | X |  |
| BCE | *M. minor* | 38.86 | -106.93 | 2,893 | X |  |  |
| NOR | *M. minor* | 39.02 | -107.07 | 3,352 | X | X |  |
| NPB | *M. minor* | 39.02 | -107.07 | 3,352 | X |  |  |
| OBJ | *M. minor* | 38.91 | -107.03 | 2,739 | X | X |  |
| CWC | *M. tilingii* | 37.59 | -118.99 | 2,784 | X | X |  |
| SON | *M. tilingii* | 38.33 | -119.64 | 2,944 | X | X |  |
| ANR | *M. guttatus* | 39.74 | -123.63 | 411 | X | X |  |
| ART | *M. guttatus* | 37.35 | -118.33 | 1,594 |  |  | X |
| BLV | *M. guttatus* | 39.01 | -107.03 | 3,311 | X |  |  |
| CCK | *M. guttatus* | 38.86 | -107.05 | 2,864 | X |  |  |
| CFC | *M. guttatus* | 38.86 | -107.17 | 2,775 | X | X |  |
| CPP | *M. guttatus* | 37.98 | -120.64 | 1,000 | X |  |  |
| DLW | *M. guttatus* | 37.07 | -118.26 | 1,249 | X |  | X |
| DWF | *M. guttatus* | 36.32 | -117.52 | 10 | X | X | X |
| GCR | *M. guttatus* | 39.49 | -106.84 | 1,402 | X | X |  |
| GJR | *M. guttatus* | 39.48 | -106.79 | 2,782 | X | X |  |
| HME | *M. guttatus* | 36.46 | -118.16 | 3,048 |  |  | X |
| JMS | *M. guttatus* | 37.77 | -119.09 | 2,651 |  |  | X |
| LLP | *M. guttatus* | 35.27 | -120.68 | 115 | X |  |  |
| LPM | *M. guttatus* | 37.63 | -119.00 | 2,594 |  |  | X |
| MBA | *M. guttatus* | 39.00 | -107.03 | 3,091 | X |  |  |
| NLC | *M. guttatus* | 37.23 | -118.63 | 3,282 |  |  | X |
| NOC | *M. guttatus* | 36.84 | -118.26 | 1,462 | X | X |  |
| NOL | *M. guttatus* | 37.23 | -118.64 | 2,981 | X | X |  |
| SAL | *M. guttatus* | 41.38 | -123.46 | 101 | X |  |  |
| SAW | *M. guttatus* | 37.20 | -119.41 | 1,208 |  |  | X |
| SMO | *M. guttatus* | 38.32 | -122.59 | 484 | X | X |  |
| TUG | *M. guttatus* | 37.95 | -120.68 | 757 | X |  |  |
| TWN | *M. guttatus* | 38.66 | -120.22 | 1,959 | X |  |  |
| ULO | *M. guttatus* | 38.88 | -106.96 | 2,801 | X |  |  |
| WFC | *M. guttatus* | 37.34 | -119.27 | 1,552 |  |  | X |
| WGL | *M. guttatus* | 38.97 | -107.04 | 3,161 | X | X |  |
| YUL | *M. guttatus* | 39.00 | -107.10 | 3,574 | X | X |  |
| **A** = Species comparison experiment; **B** = Elevation-of-origin experiment; **C** = Eastern Sierra Nevada experiment. | | | | | | | |

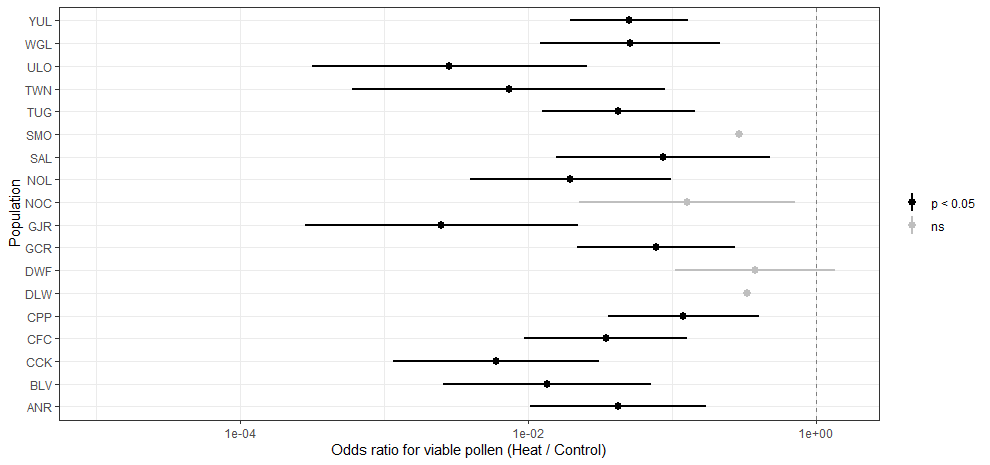

Figure S1. Odds ratio in the species comparison experiment show some populations had non-significant differences in pollen viability between heat and control. Populations with extremely imprecise estimates were omitted from the figure for visualization because their confidence intervals were too wide to display clearly.

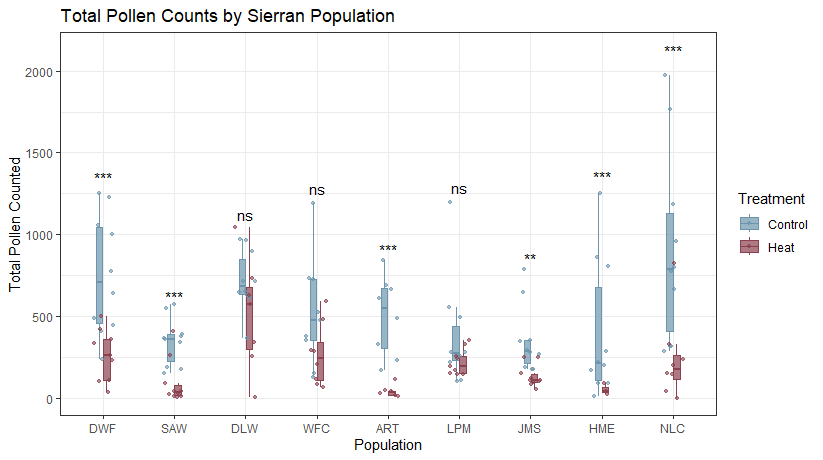

Figure S2 Total pollen counts varied in eastern Sierra Nevada populations. Asterisks indicate significant difference between control and heat treatments within population comparisons.
